# *In vivo* optical clearing of the mouse brain

**DOI:** 10.64898/2026.08.18.745591

**Authors:** Maiko Kume, Ningdong Kang, Alejandro Akrouh, Dae Woo Kim, Joshua T Dearborn, David F Wozniak, Daniel Kerschensteiner, Timothy E. Holy

## Abstract

Light microscopy is one of the most powerful tools for understanding living systems, but the opacity of tissue prevents visualization of all but superficial layers. Several methods to clarify tissue have been developed, but most require fixed specimens. To address the challenge of improving resolution in functioning neuronal circuits, we developed a biocompatible clearing agent, iodixanol-ACSF, which is capable of increasing the transparency of living neuronal tissue. Brain-cleared mice were motile and unimpaired on a variety of behavioral tasks, and extracellular recordings showed that many cellular and circuit phenomena were well-preserved. In live iodixanol-ACSF cleared mouse brain tissue, both transmission and cellular-resolution fluorescence microscopy indicate improvements of 150-200% in penetration depth with one-third to one-half the laser intensity when compared to untreated tissue. Our results show that iodixanol-ACSF clearing will enable deeper imaging and extend our understanding of neuronal circuit function.

## Introduction

Light microscopy has led to numerous breakthrough discoveries in the biological sciences. However, a persistent limitation to optical imaging is the degradation of image quality at increasing depths. The main sources of degradation are absorption, aberration, and scattering. Scattering has been shown to contribute more to the reduction of light transmission than absorption at the visible and near-infrared wavelengths used for fluorescence microscopy^1^. Adaptive optics can address sample-induced, bulk aberrations which are spatially and temporally stable^2,3^. However, adaptive optics cannot correct for scattering^4^. Scattering is a result of the fine-scale heterogeneity of refractive indices within the sample arising from mismatches between the extracellular fluid (ECF) and wavelength-scale cellular components^5^. ECF has a refractive index of 1.34, similar to cerebral spinal fluid (CSF), while intracellular components have higher refractive index values (cortical cytoplasm = 1.353-1.368, mitochondria = 1.38-1.41, nucleus = 1.39-1.43)^6–8^. In neural tissue, the many fine membrane-bounded neurites are an additional contributor to scattering.

In fixed tissues, there are numerous techniques for increasing transparency^9–25^. While use of such clearing techniques has provided unprecedented optical access of the brain, fixation and the harsh processing steps such as dehydration, lipid extraction and rehydration of tissue with a solution of higher refractive index are incompatible with living tissue. One previous *in vivo* technique may provide modest improvement, but it can only be seen in group averages^26^. A more recent method has not been shown to be compatible with brain imaging^27^.

Here we describe an approach that reduces scattering by raising the refractive index of ECF while still maintaining neural health and function. We designed a custom high-refractive index isoosmotic artificial cerebrospinal fluid (iodixanol-ACSF or I-ACSF) that preserves much of the ionic balance necessary for normal neuronal function. We first demonstrate that I-ACSF is capable of significant clearing in brain slices when viewed by transmission microscopy. To examine its effects on neuronal function, we continuously perfused clearing agent into the ventricles while the mice underwent a battery of behavioral tests. To further characterize any perturbations of activity within a neuronal circuit, we performed comparative multi-electrode array recordings of the retina in control and I-ACSF solutions. Finally, to demonstrate the utility of this technique for neuronal imaging, we characterized I-ACSF’s improvements in resolution and imaging depth of fluorescently-labelled neurons using confocal and two photon microscopy. We thus demonstrate a straightforward method of altering the optical properties of living neuronal tissue that reduces scattering and improves image quality. We anticipate that *in vivo* clearing can push us closer to bridging the gap between neurophysiology and animal behavior.

## Results

### Evaluation of tissue clearing capability

To obtain the greatest increase in transparency for a given concentration of clearing agent, we focused on biocompatible organic compounds containing a large number of heavy ions, as the inner-shell electrons endow the material with a large dipole moment. These characteristics are satisfied by radiocontrast agents^5,19,28,29^. We focused on iodixanol, a non-ionic dimer of iodinated benzenes that can be formulated to be isotonic to plasma^30^. Iodixanol is clinically-approved for human use and is routinely used for X-ray based visualization of vasculature. Iodinated contrast agents have been used previously to modulate refractive index. For example, contrast agents have been added to immersion media to match its refractive index to that of *ex vivo* cleared samples (SeeDB2, ACT-PRESTO, SWITCH). More recently, iodixanol has been used as a non-toxic refractive index matching fluid for live specimens such as planaria and zebrafish embryos^31^. Encouragingly, Boothe et al. showed that iodixanol did not have a deleterious effect on the growth and survival of these organisms at concentrations relevant to physiological refractive index matching. However, all of these efforts were directed at reducing the bulk mismatch in refractive index between the sample and the exterior immersion media, in much the same way that oil immersion fluid matches the refractive index of glass coverslips.

Here, our intent is to correct for finer inhomogeneities in refractive index within a sample that lead to scattering and cannot be easily corrected using tools such as adaptive optics. This requires index-matching in the extracellular fluid throughout tissue, for which we designed I-ACSF. The composition I-ACSF (see Methods) was designed to provide a high refractive index of 1.385 while satisfying the conflicting demands of maintaining an osmolarity matching that of normal ACSF (310 mOsm), an ionic milieu that perturbs Nernst potentials by no more than 10mV, and avoidance of hyponatremia.

We first tested whether I-ACSF increases optical clarity by measuring its effect on tissue transmittance. We immersed 200 µm thick slices from the mouse brain in control artificial CSF (ACSF, n = 1.34) or I-ACSF. We then overlaid the slices on an opaque edge of a 1951 USAF resolution target to image the transmission through the slice. On a qualitative, macroscopic scale, we observed that finer patterns were visible through the I-ACSF treated slices while not resolvable in equivalent thickness slices in ACSF (Figure 1a). 10 µm sized features were easily resolved through 200 µm of I-ACSF treated tissue, while we struggled to resolve features double this size through ACSF-treated tissue (Figure 1b). Only heavily myelinated structures such as the corpus callosum and internal capsule failed to show substantial clearing, which is expected since I-ACSF is aqueous and the refractive index of the lipid-rich myelin is higher than that of gray matter^32^. When we increased the concentration of iodixanol without attempting to maintain physiological osmolarity to obtain a refractive index of 1.42, there was an increase in transparency in myelinated regions but at a cost of tissue expansion, highlighting the tradeoff between transparency and maintaining physiologically relevant parameters (**Supplemental Figure 1a**). Transmission through a slice thickness of 200 µm is a proxy for the total optical path length encountered during epi-illumination at depths of 100 µm (Figure 1c), which is approximately the limit for high-quality imaging by single-photon techniques^33,34^.

**Figure 1.**
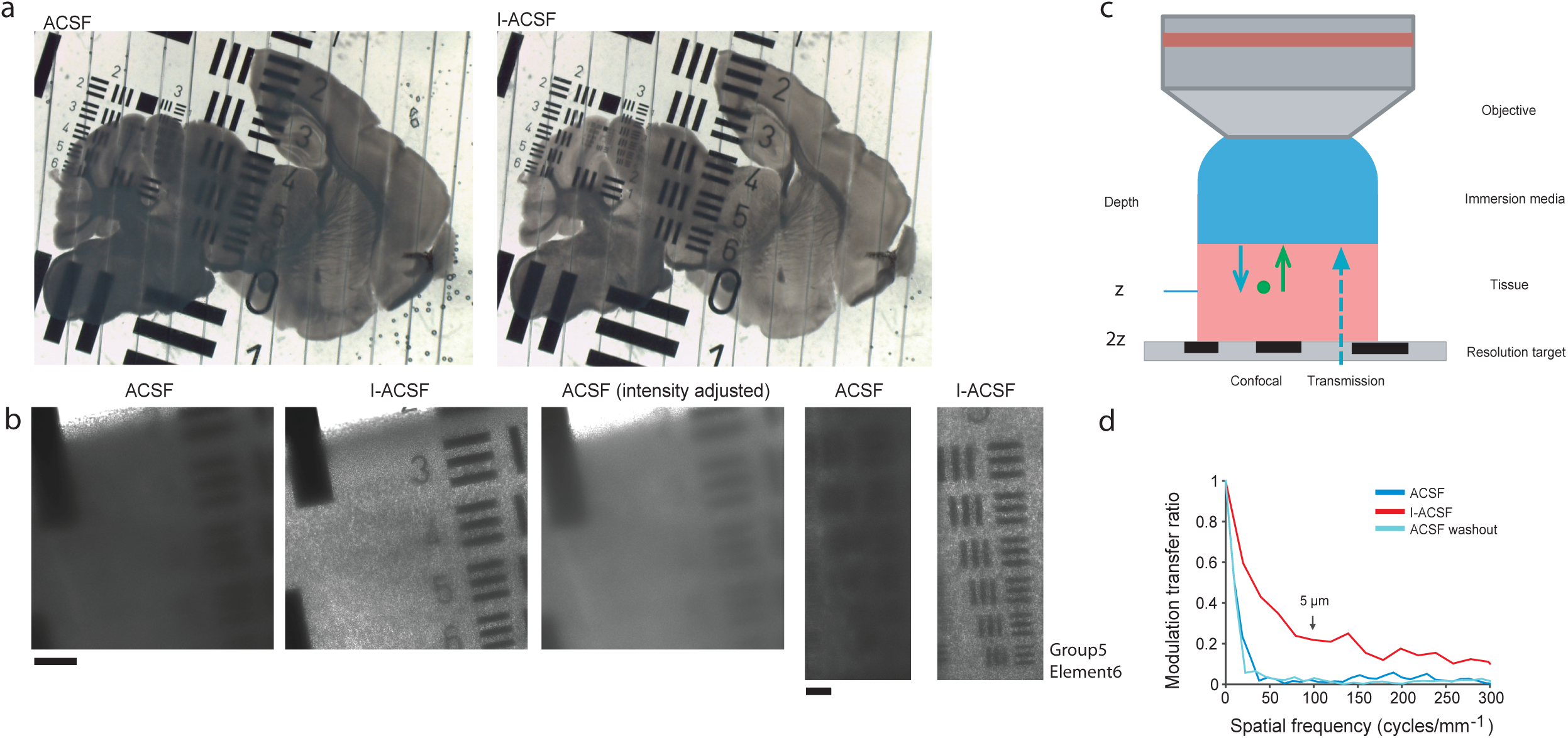
Clearing of brain slices in transmission optical imaging. (**a**) Bright field image of a resolution target imaged through 200 µm of brain tissue following incubation in ACSF, then 30 minute incubation in I-ACSF. (**b**) Confocal transmission images of a single 200 µm thick slice, imaged in ACSF (left) and I-ACSF (center). 20 µm features (group 4, element 5 USAF resolution target) were clearly visible through the tissue imaged in I-ACSF. When the mean intensity of the ACSF image was increased to match the mean intensity of the I-ACSF image, the element numbers (3-6) still could not be read through the tissue. Scale bar represents 100 µm (left three panels). 8.8 µm features from group 5 element 6 of the resolution target were clearly resolved through 200 µm of I-ACSF incubated tissue but not through ACSF incubated tissue. Scale bar represents 50 µm (right two panels). (**c**) A representation of the optical path through tissue in confocal and transmission imaging configurations (blue represents illumination, green emitted fluorescence) (**d**) The modulation transfer function calculated from confocal transmission images of the resolution target through 200 µm of cortical tissue.

To quantify the improvement in clarity, we calculated the modulation transfer function (MTF), which measures the degradation of image quality for both coarse- and fine-scale features. The transfer function rapidly dropped below 5% of its initial value at a feature size corresponding to 12.5 µm (40 lp/mm), indicating a loss of cellular resolution once a photon travels through 200 µm of tissue (Figure 1d). On the other hand, the MTF measured through the I-ACSF treated tissue had a more gradual drop-off and at 100 lp/mm (equivalent to 5 µm feature), the curve remained above 20% of the initial value. These values approximately correspond to what we observed visually with the resolution target (Figure 1a,b). We also observed that the clearing effect can be reversed by washout with standard ACSF (Figure 1d).

To separate the effect of iodixanol clearing on scattering from its effect on aberrations, we also imaged the tissue with a low numerical aperture (NA) objective (NA = 0.082). The larger angle light rays accepted by a high NA objective are refracted more than those closer to the optical axis. By using a low NA objective, we reduced the impact of aberrations. We found that the MTF curve drop-off was always more gradual for the cleared tissue, irrespective of objective parameter or imaging system, demonstrating that iodixanol treatment reduced scattering (**Supplementary Figure 1b**). We conclude that I-ACSF treatment increases light transmission through tissue and has potential as an *in vivo* clearing agent.

### Behavioral testing of iodixanol treated mice

One potential concern with *in vivo* clearing is whether it perturbs neuronal function. To test for large-scale impacts on sensory, motor, and/or cognitive function, mice receiving a brain-wide infusion of iodixanol through the ventricular system were subjected to a battery of behavioral tests. We compared the performance of control mice infused with ACSF with those infused with Visipaque 320, a formulation of iodixanol designed to be isotonic to blood plasma (n_ACSF_ = 10, n_Iodixanol_ = 12). This characteristic of Visipaque enabled us to deliver it directly into the brain while minimizing the risks associated large osmotic imbalances^8^. The higher iodixanol concentration in Visipaque (420 mM) compared to concentration in I-ACSF (184 mM) served as a conservative upper bound on any behavioral abnormalities from ventricular infusion of this compound.

Using a cannula placed into the right lateral ventricle and an implanted osmotic pump, Visipaque was continually delivered at a rate approximately equivalent to 40% of the total CSF production rate in mice^35^. To reduce the dilution of iodixanol by CSF and the risk of excess intracranial pressure, we co-infused with acetazolamide (AZ), a carbonic anhydrase inhibitor shown to reduce CSF production by up to 60%^36^. After approximately 24 hours of infusion, both groups of mice were still mobile, eating, drinking and defecating and appeared to be in good general health (**Supplementary video 1a,b**). At this point, exploiting iodixanol’s use as a radiocontrast agent, we imaged the mice using X-ray computed tomography (CT), to confirm the presence of iodixanol in the lateral ventricles in each mouse used for behavioral testing (see below). To ensure equal treatment, mice receiving ACSF-AZ were also imaged.

After confirming infusion, following CT imaging and at least 30 minutes after the end of isoflurane exposure, the same groups of mice underwent a battery of behavioral tests performed by an observer blinded to the contents of the pump. Intact, non-infused mice were tested in addition to observe any change in behavior related to the surgery, the implantation of the cannula and the pump, or the ventricular infusion (n_control_ = 10). The infused mice were tested with the cannula and osmotic pump in place. The mice were tested on locomotor activity/exploratory behavior, sensorimotor measures, and olfactory preference, using a set of tasks selected to be thorough yet accomplishable within a single day of testing. All mice survived the infusion, CT imaging and behavioral testing.

Unsurprisingly, the results from the behavioral tests revealed some differences between unmanipulated animals and infused animals. We also noted further surgery-independent differences between ACSF-infused mice and iodixanol-infused mice; these differences generally depended on whether the task was “motivated” or “spontaneous.”

For assays of “motivated” activity, the presence or absence of clearing agent was less important than the effect of surgery (**Figure 2a**). No differences in performance were detected between ACSF-infused and iodixanol-infused mice among 9 sensorimotor measures. In 7 of 9 of these same measurements, no differences were observed in comparisons between non-surgery and iodixanol-infused mice. Iodixanol-infused mice performed comparably to the non-surgery mice on time spent on a ledge, pole climb down time, turning and climbing down a pole, and time hanging on to 60-degree, 90-degree and inverted screens. ACSF-infused mice also performed comparably to non-surgery mice in 7 of 9 sensorimotor assays. Differences between the non-surgery mice and the infused mice were observed in time on an elevated platform (p_C-A_ = 0.0012 and p_C-I_ = 0.0007), time to climb a 90-degree inclined screen (p_C-A_ = 0.00005 and p_C-I_ = 0.0008), time to climb a 60-degree inclined screen (p_C-I_ = 0.001), and time hanging upside down on an inverted screen (p_C-A_ = 0.01), indicating some effect on balance, coordination, speed of movement, and strength due to surgery, pump implantation, and/or fluid infusion.

**Figure 2.**
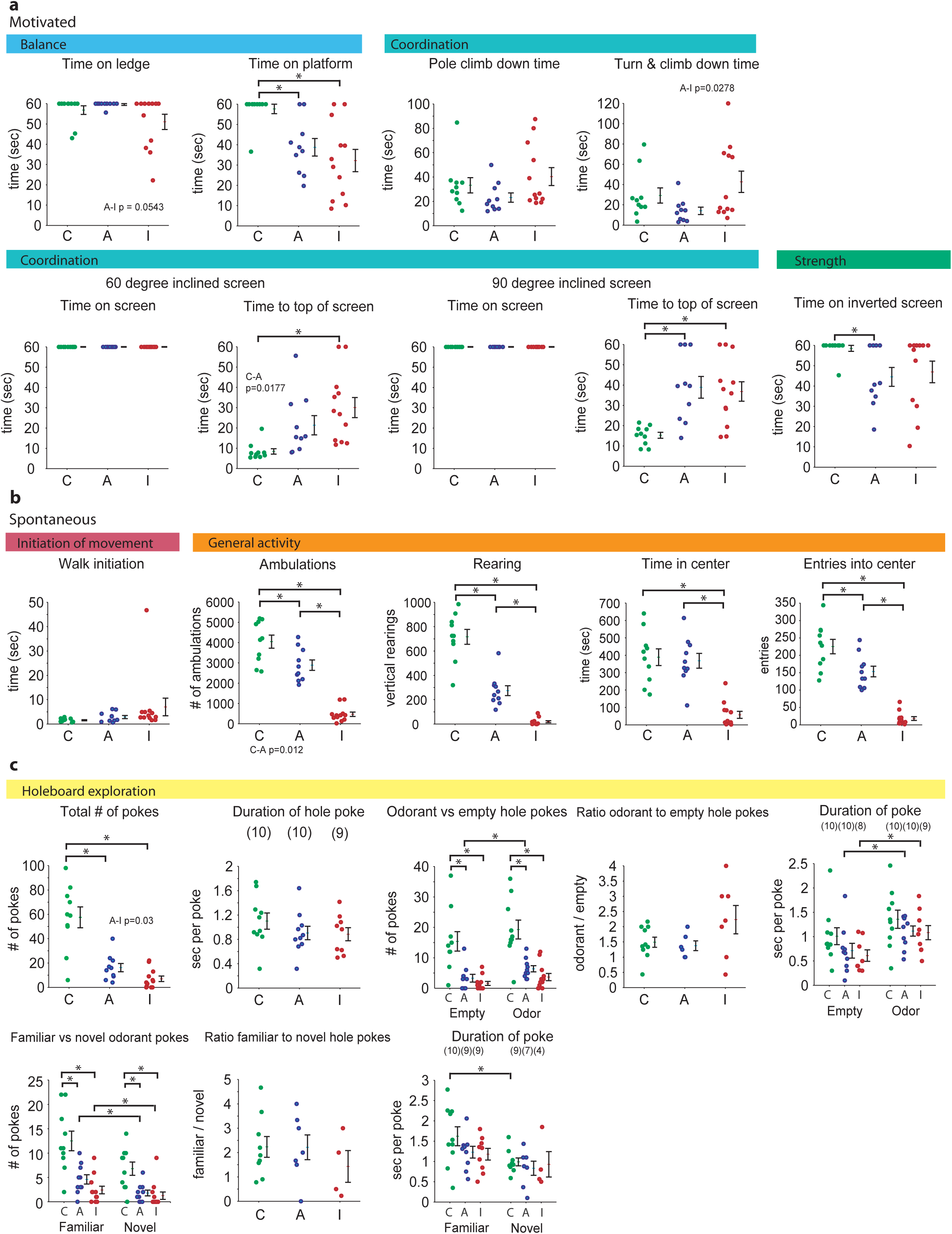
Behavioral testing of mice with intraventricular infusion of iodixanol. (**a**) Mice receiving an infusion of iodixanol and acetazolamide (I, n = 12), mice receiving an infusion of ACSF and acetazolamide (A, n = 10), and control mice with no manipulation (C, n = 10) were evaluated in terms of their performance on a battery of behavioral assays. The infused mice did not differ among the two groups in any of the sensorimotor tests involving balance, coordination, and strength. Both infused groups performed worse than the control group with regards to remaining on an elevated platform (p_C-A_ = 0.0012 and p_C-I_ = 0.0007 and climbing to the top of a vertical screen (p_C-A_ = 0.00005 and p_C-I_ = 0.0008). The I-ACSF infused group but not the ACSF infused group took a significantly longer time than the control group to climb to the top of a 60 degree inclined screen (p_C-I_ = 0.001). The ACSF infused group spent significantly less time than the control group hanging upside down on an inverted screen (p_C-A_ = 0.01). (**b**) Differences were observed between the groups on assays of spontaneous activity. While the times taken to move out of a restricted space (walk initiation) did not differ among the groups, each group differed from the other in the number of total ambulations, i.e., whole body movements (p_C-A_ = 0.0120, p_C-I_ = 4×10^-11^ and p_A-I_ = 10^-8^) and in the number of vertical rearings (p_C-A_ = 10^-5^, p_C-I_ = 7×10^-11^ and p_A-I_ = 10^-6^). The entries into the center, time spent in the center, and distance traveled in the center, which may represent a combination of locomotor activity and discomfort in the open field, differed between ACSF and I-ACSF infused groups and between the control and the I-ACSF groups. The number of entries in the center differed between the control and ACSF groups. (**c**) The number of total hole board exploration pokes did not differ between the ACSF and I-ACSF groups. However, both groups made significantly fewer hole pokes than the control group (p_C-A_ = 0.0003 and p_C-I_ = 5×10^-6^). The decrease in hole poke frequencies among both the infused groups compared to the control group was found regardless of the contents of the hole (empty, odorant, familiar or novel). No differences were detected in the durations of hole pokes between the groups.

“Spontaneous” activity measures were reduced in the both groups of infused mice compared to the non-surgery control; however, this effect was greater in the iodixanol-infused mice (**Figure 2b**). When examining such “spontaneous” behavior, no differences were observed between groups in time to initiate movement out of a circumscribed space. However, following a thirty-minute locomotor activity test, ACSF-infused mice had less ambulatory activity (p_C-A_ = 0.0120), fewer rearings (p_C-A_ = 10^-5^) and fewer entries into the center of the field (p_C-A_ = 0.0121) compared to unmanipulated mice. Iodixanol-infused mice had additional reductions in activity in these categories. Total ambulations (p_C-I_ = 4×10^-11^ and p_A-I_ = 10^-8^), number of rearings (p_C-I_ = 7×10^-11^ and p_A-I_ = 10^-6^), time spent in the center (p_C-I_ = 10^-6^ and p_A-I_ = 9×10^-7^), entries into the center (p_C-I_ = 10^-9^ and p_A-I_ = 2×10^-8^) and distance traveled in the center (p_C-I_ = 4×10^-9^ and p_A-I_ = 2×10^-8^) were reduced in iodixanol-infused mice compared to both unmanipulated and ACSF-infused mice. Overall, in empty cages the iodixanol-infused mice spent less time moving than the other groups.

Another spontaneous task, exploration of a hole-board, likewise revealed a substantial effect of the surgery, but the differences between ACSF and iodixanol-infused mice were smaller or nonexistent (**Figure 2c**). Total hole poking frequency was reduced in both infused groups compared to the non-infused mice (p_C-A_ = 0.0003, p_C-I_ = 5×10^-6^ and p_A-I_ = 0.030, the latter not significant when corrected for multiple comparisons), but no difference was found in hole poke duration. Altogether, these results suggest few differences for “motivated” behaviors but show evidence for limited iodixanol-mediated decrease in “spontaneous” behaviors such as exploration of the test cambers. Crucially, despite the manipulations, all animals were alive, were alert, and were capable of performing all of these behavioral tasks.

After behavioral testing, we quantified brain concentrations of iodixanol-infused mice by calibrating our CT images (**Figure 3a**) with a set of iodine standards (**Figure 3b,c**). The average concentration of iodixanol delivered into the brain tissue (excluding the Visipaque-filled ventricles) after 24 hours was 5.6 mM of iodixanol, which can be estimated to be equivalent to 28 mM of iodixanol in ECF. This estimate assumes that iodixanol does not enter the cell, and 20% of cortical volume consists of extracellular space^37,38^. Since we observed variability among iodixanol concentration and behavioral results, we examined whether performance could be explained by iodixanol concentration. We found no correlation between the concentration of iodixanol measured in the brain and performance on any behavioral test (p > 0.05, **Figure 3d** and **Supplementary Figure 2**).

**Figure 3.**
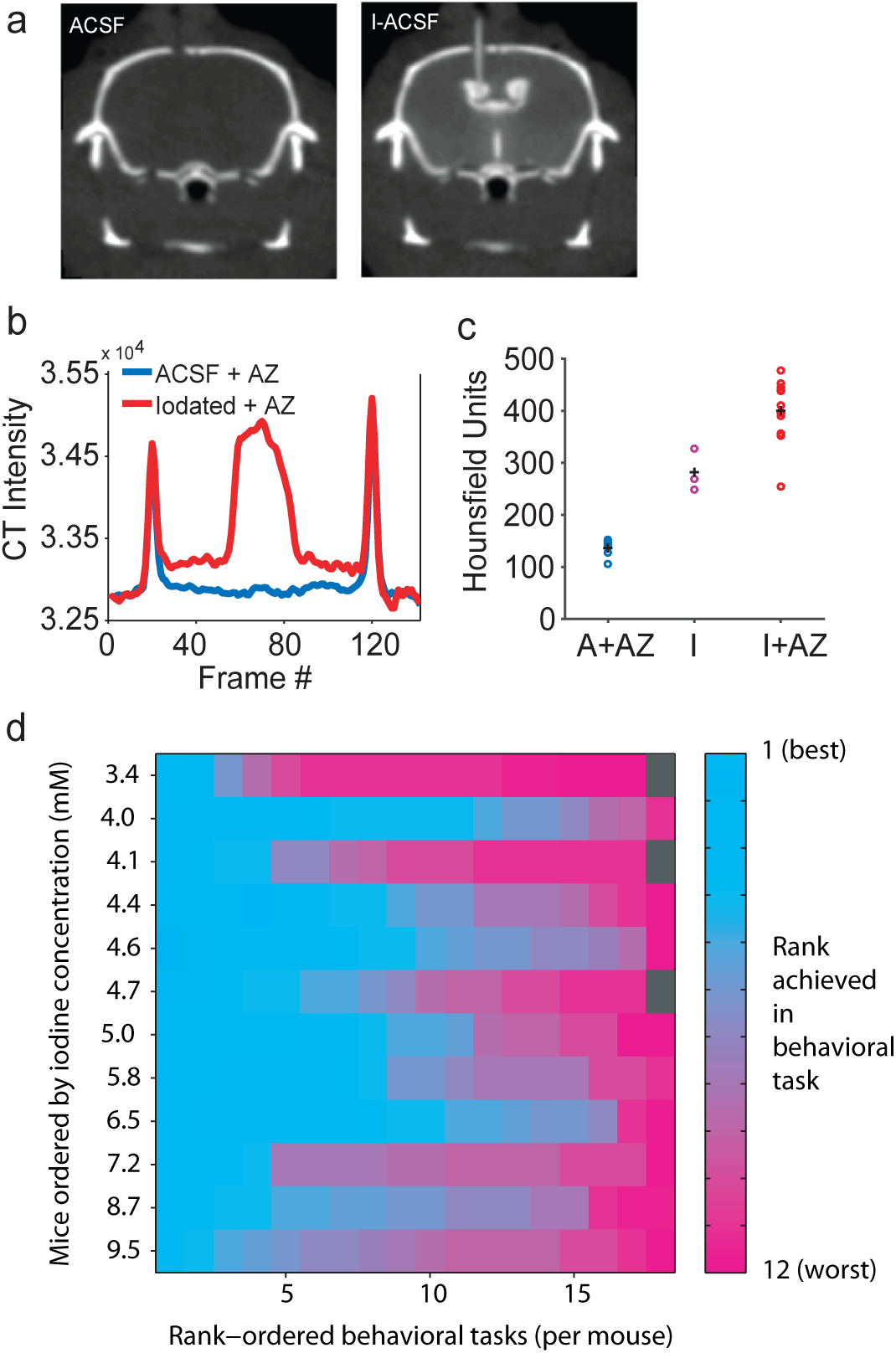
Whole brain delivery of iodixanol through the lateral ventricles. (**a**) Coronal CT image sections of ACSF and iodixanol-infused mice, with the lateral ventricle and cannula visible only in the latter. (**b**) A sample line-section taken through the skull and lateral ventricles of ACSF and iodixanol treated mice. (**c**) Radiodensity of brain tissue in ACSF and iodixanol-infused mice after 24 hour infusion. (**d**) The relative performance of the iodixanol-infused mice at each behavioral task compared to the concentration of iodine measured in brain tissue. Each row represents a single mouse. The order in which tasks are displayed varies among rows. The tasks are arranged in order of performance for each test among the mice. Rank is assigned based on difference in performance from the non-manipulated mice. Gray shading indicates that the mouse did not perform the task.

### Evaluate effect on circuit physiology

To further characterize the effects of clearing on neuronal function, we performed extracellular spike recordings from an intact circuit. Extensive work in the visual system has shown that the developing retina generates spontaneous activity patterns (i.e., retinal waves), which instruct circuit refinement downstream^39^. The spatiotemporal properties of retinal waves have been well-characterized and shown to be highly sensitive to pharmacological perturbation. Using multielectrode array recordings, we compared spontaneous activity in the postnatal day 5 to 7 ganglion cell populations in ACSF and I-ACSF solutions. Most importantly, waves involving the firing of large fractions of ganglion cells on the array persisted in I-ACSF (**Figure 4a,b**). Furthermore, although the frequency of retinal waves was reduced in I-ACSF (**Figure 4d**, P = 0.002), overall firing rates of ganglion cells (**Figure 4c**, P = 0.3), their firing rates during waves (**Figure 4e**, P = 0.4), and spatiotemporal patterns of activity propagation (**Figure 4f**, P = 0.7) were unchanged. By contrast, the commonly used anesthetic isoflurane (2%) suppressed most ganglion cell firing (**Figure 4g-i**, P = 0.04) and abolished retinal waves (**Figure 4h**, P = 0.04). Thus, iodixanol has comparatively mild effects on network activity patterns in a developing circuit.

**Figure 4.**
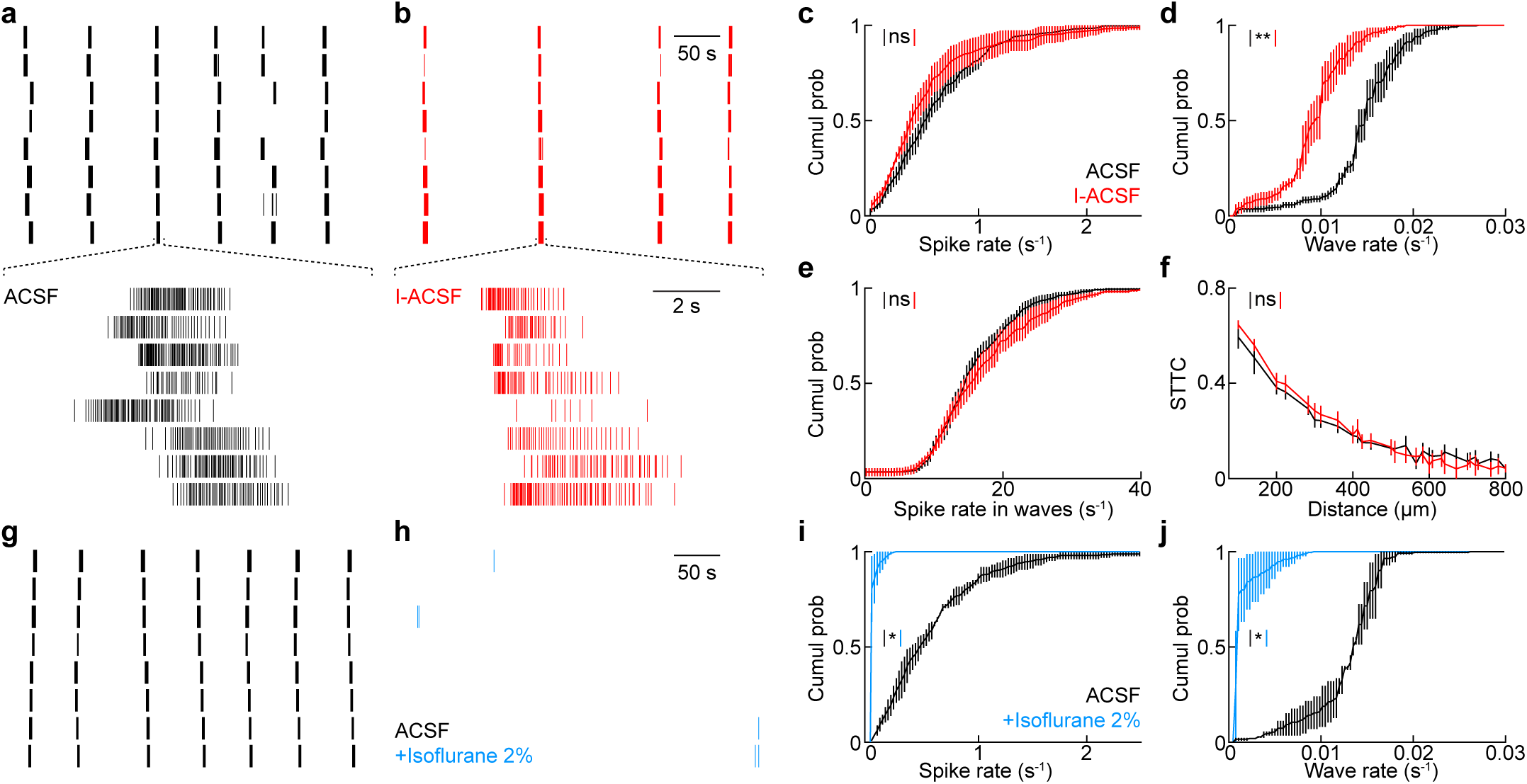
Patterned spontaneous activity persists in I-ACSF. **a, b**, Representative spike rasters of eight ganglion cells (one per row) in ACSF (**a**, black) and I-ACSF (**b**, red) show correlated bursts of activity (i.e., retinal waves). **c-e,** Cumulative distributions of ganglion cell firing rates (**c**), rates of retinal waves (**d**) and ganglion cell firing rates within retinal waves (**d**) in ACSF (black) and I-ACSF (red, n = 272 cells, n = 6 retinas). **f,** Summary data of spike time tiling coefficients (STTC) as a function of the distance between pairs of recorded cells (n = 6,390 pairs, n = 6 retinas). **g,h,** Representative ganglion cell spike rasters in ACSF (**g**, black) and ACSF with 2% isoflurane (**h**, blue). **i,j,** Cumulative distributions of ganglion cell firing rates (**g**) and rates of retinal waves (**h**) in ACSF (black) and ACSF with 2% isoflurane (blue, n = 212 cells, n = 2 retinas). Throughout the figure, ** indicates P < 0.01 and ns indicates no significant differences for statistical comparisons.

### Single photon fluorescence imaging of cleared living neuronal tissue

Multiphoton microscopy has been widely embraced for imaging biological tissue due to the ability to image with reduced scattering. However, single photon imaging techniques such as light sheet microscopy and light field microscopy have different strengths compared to point scanning approaches used by standard two photon microscopes. Many super resolution imaging techniques are single photon techniques as well, and such techniques could benefit from a reduction in scattering. To characterize improvements in fluorescence image quality due to clearing, we imaged living tissue slices with GFP-labelled neurons with confocal microscopy, a single photon imaging technique which ordinarily experiences severe depth limitations due to scattering. Since slices are mostly flat, they are less susceptible to bulk aberrations compared to a curved surface like the brain. In addition, slices do not have the outward flux of CSF counteracting the diffusion of iodixanol as in a live mouse. Since clearing performance depends on local tissue properties, for the most direct comparison, we imaged the same region of the same tissue in ACSF and I-ACSF by gradually replacing the media without perturbing the tissue while maintaining identical imaging parameters. To accommodate the unconventional refractive index (1.385) of the immersion solution, we used an off-the-shelf, variable immersion objective (HCPLAPO IMM CORR CS 20x NA 0.7 WD 0.26, Leica) and determined the correction collar setting with the best performance in our clearing media by measuring the point spread function of sub-diffraction fluorescent beads (**Supplementary Figure 3a** and **Methods**). Once the correction collar was set, GABAergic interneurons in a GAD65-GFP mouse^40^ were imaged in coronal slices, focusing on the cortical region.

We were able to achieve sub-cellular resolution substantially deeper in cleared tissue than in untreated tissue (**Supplementary Figure 3b,c**). When imaging at 110 µm depth, the same cell bodies were over ten-fold brighter, and submicron-sized processes could be visualized only when imaged with I-ACSF superfusion (**Supplementary Figure 3d**). Clearing as a function of time was monitored in both ACSF and I-ACSF. Clearing was not observed in ACSF-incubated tissue at longer incubation time, indicating that the process was not due to change in tissue health over time. Reduced scattering had a large impact on image intensity, as the laser power could be reduced by a half to two-thirds in I-ACSF compared to ACSF, indicating more efficient transport of photons into and out of the tissue (**Supplementary Figure 3d)**.

### Two photon microscopy of the cleared living mouse brain

Two photon microscopy has been invaluable for performing *in vivo* imaging and allowing access to deeper regions of the brain. Two photon microscopy enables deeper imaging by utilizing the closely timed arrival of two photons of approximately double the typical excitation wavelength for a fluorophore. Longer wavelengths undergo less scattering, and when used in point-scanning the emitted photons do not need to be imaged since the large majority of collected photons can be attributed to the objective’s focal spot. However, the loss of intensity caused by the attenuation of light from scattering is still the limiting factor for achievable depth in two photon microscopy^4,41^.

To investigate the utility of iodixanol clearing for *in vivo* applications, we first performed structural imaging in an anesthetized mouse. To reduce the effect of experimental variability (including damage from microinjection), the mouse was head-fixed under the microscope for imaging while either ACSF or I-ACSF was superfused over the exposed brain, and the same brain region was imaged before and after treatment. While maintaining identical imaging parameters across the two conditions, we observed that the same GFP labelled neurons were dramatically brighter after superfusion with I-ACSF (**Figure 5a**). The signal to background ratio of the neurons increased by two to three-fold after two hours of I-ACSF superfusion (**Figure 5b**). We could image neurons deeper after I-ACSF treatment at laser powers of 25 mW, measured at the front of the objective (**Figure 5c**). The clearing effect increased over the course of two hours (**Supplementary Figure 4a**), indicating that the increase in signal intensity cannot be attributed solely to a correction of bulk aberrations. These changes were not observed after superfusion with only ACSF for similar durations, indicating that the clearing effect cannot be attributed to superfusion time (**Supplementary Figure 4b**). The clearing effect was reversible and repeatable on the same mouse if the window was sealed between imaging sessions.

**Figure 5.**
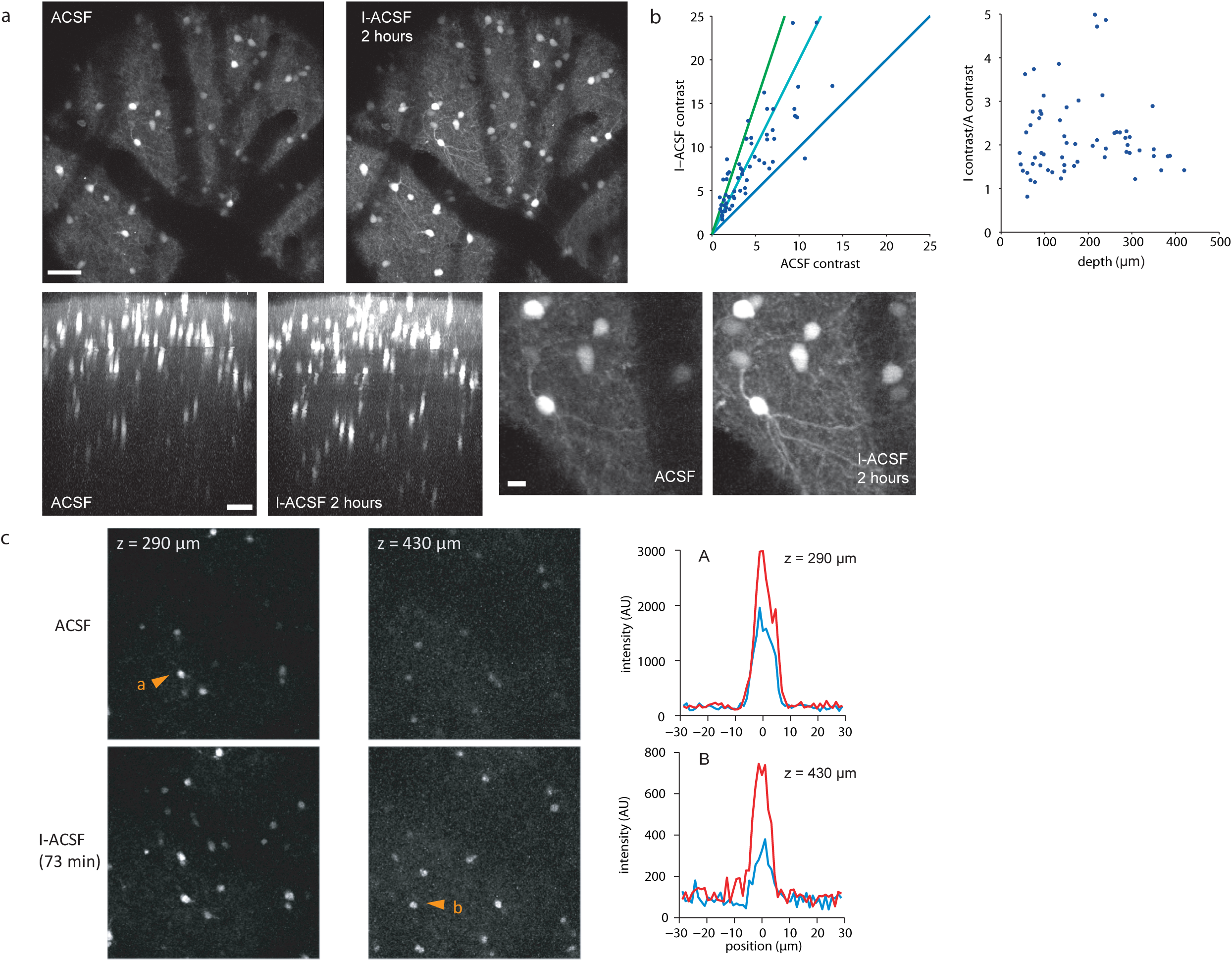
Two photon imaging of the cleared live mouse brain. (**a**) Maximum intensity projection images (axial and lateral) of GFP positive neurons in the mouse cortex before and after I-ACSF superfusion. The mouse was awake during imaging. Images are displayed as the square root of the original intensity to condense dynamic range. Zoomed in view of same region, showing the improvement in resolution of fine scale features. Scale bar indicates 50 µm and 10 µm in the zoomed in image. (**b**) Plotting the contrast of the same neurons in (a) in each condition shows the improvement in resolvability of cells. (**c**) Individual neurons, before and after I-ACSF superfusion. The mouse was anesthetized during imaging. The intensity plots of cross sections of neurons shows an improvement in resolvability in I-ACSF.

Next, we proceeded to examine whether iodixanol clearing can be used with calcium imaging. Imaging was performed on mice with genetically encoded GCaMP6f expression in excitatory neurons. The mice were also injected with AAV1.CAG.tdTomato.WPRE.SV40 to provide a fixed anatomical reference across conditions. To stabilize the brain against movement while allowing rigorous quantification via before/after comparisons, we replaced the conventional glass coverslip with a circular piece of titanium mesh that permitted both imaging and direct fluid access to the brain (**Figure 6a**). After surgery, we imaged mice with ACSF or I-ACSF superfusion. We observed small amounts of lateral movement in the mice, which could be corrected with cross correlation-based registration. To characterize any effect of iodixanol treatment on neuronal function, we compared activity during superfusion with ACSF and I-ACSF.

**Figure 6.**
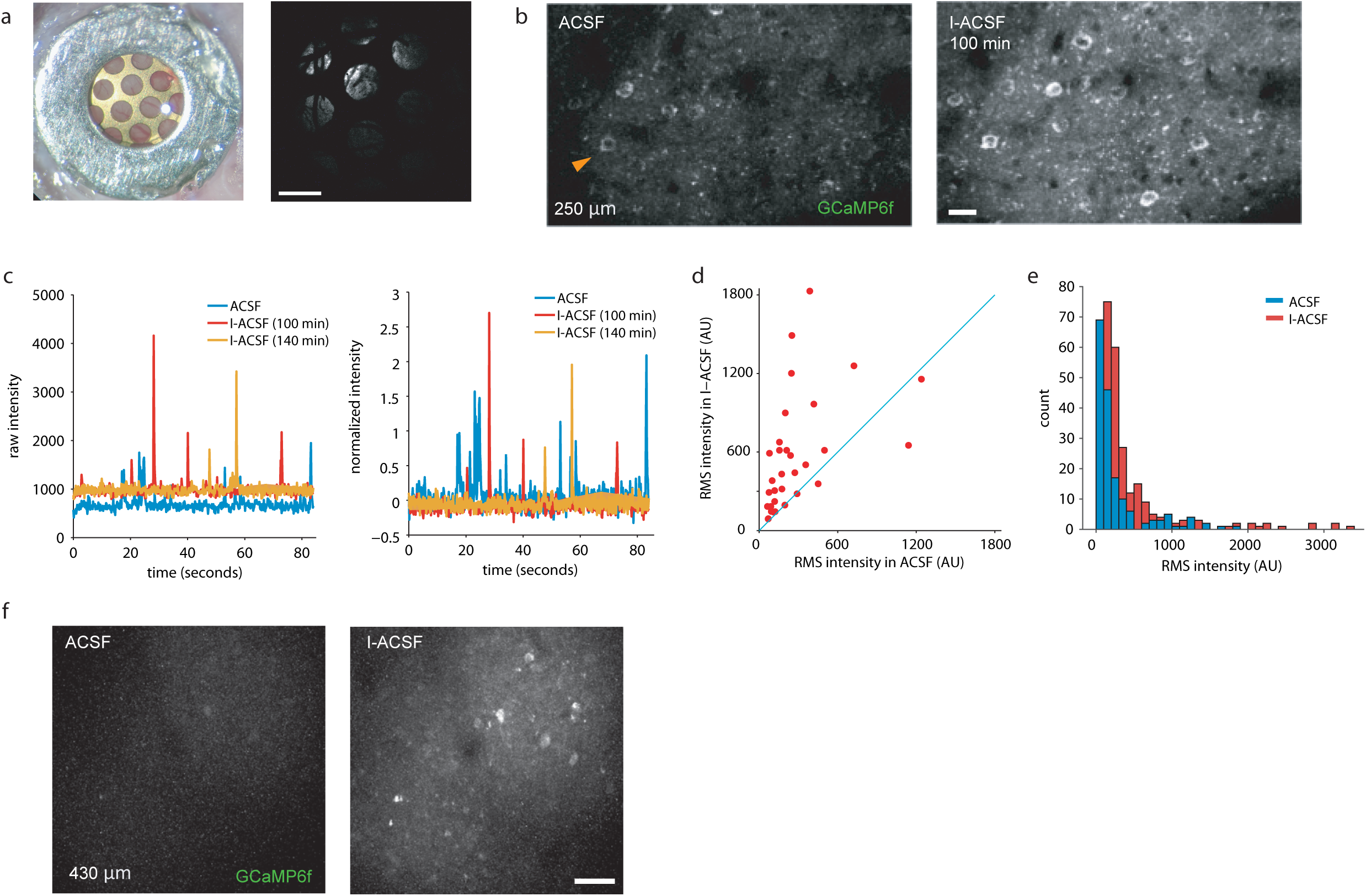
Two photon imaging of calcium activity in the cleared live mouse brain. (**a**) The titanium mesh was attached to cranial window and provided rigidity and simultaneous optical and fluid access to the brain. Scale bar indicates 500 µm. (**b**) Two photon imaging performed on GCaMP6f and tdTomato labelled neurons, imaged through the titanium mesh window. Scale bar indicates 20 µm. (**c**) Traces of calcium activity in neurons imaged during superfusion with I-ACSF. The raw intensity and delta F/F traces are shown for the cell marked in b). (**d**) The RMS intensity of neurons active in both imaging conditions matched using CMNF. (**e**) The RMS intensities of all identified cells from imaged under both superfusion conditions (n = 4 mice). (**f**) GcaMP6 positive cells can be detected at a depth of 430 µm under I-ACSF superfusion while visualization was challenging under ACSF superfusion with identical imaging parameters. Scale bar indicates 50 µm.

Neuronal activity persisted in the presence of iodixanol treatment (**Figure 6b,c, Supplemental Video 2a,b**). The same neurons could be identified and imaged with higher mean fluorescence intensity after superfusion with I-ACSF (**Figure 6d**). Under the same laser intensity that allowed imaging at 430 µm depth during I-ACSF superfusion, most of the same neurons could not be visualized during ACSF superfusion **(Figure 6e**). Calcium activity could continue to be observed at time points exceeding three hours after start of I-ACSF superfusion. We conclude that I-ACSF provides dramatic enhancement in image quality and intensity in living, functional neuronal tissue.

## Discussion

Much of our understanding of the brain relies on observing neurons in their intact, functional state. Here, we described a clearing method that enables improved imaging resolution and depth in the intact mouse brain without compromising the ability to record activity. Chemically manipulating the properties of live tissue is a balancing act between keeping the tissue healthy and achieving the desired effect, in our case, introducing a high concentration of an extraneous compound to optically clear tissue. Our technique is not the first attempt at clearing live tissue. Topical application or dermal injection of hyperosmotic sucrose, glycerol and polyethylene glycol solutions have been shown to increase skin transparency for imaging blood flow^42–45^. Similarly, a technique to clear the skull enabled the imaging of underlying cortical vasculature and fluorescently labelled dendrites and microglia^46,47^. However, none of these preserves the ionic and osmotic conditions needed for healthy neuronal tissue.

To test our method while reducing preparation-to-preparation variability, we chose to perform acute imaging with direct I-ACSF superfusion. We found that diffusion can deliver enough I-ACSF to reduce refractive index mismatch and quantitatively reduce scattering within fresh tissue. However, we have no reason to believe that superfusion is special, and other delivery methods may prove equal or superior for regular use.

In addition to enhanced spatial resolution, a major advantage of our technique is the improved brightness and/or ability to reduce laser power. In living samples, minimizing total photon dose is the primary means to reduce phototoxicity and thermal damage^48^. Ultimately, the ability to tune the refractive index to reduce scattering *in vivo* is applicable to many other optical imaging techniques since scattering is pervasive and difficult to correct computationally.

While this manuscript was in initial preparation, an alternative approach to *in vivo* clearing was reported^26^. This alternative approach, called MAGICAL, involves supplementing the drinking water of the mouse with glycerol. While this mode of delivery has the benefit of simplicity, the size of the clearing effect is much smaller than we report here. With MAGICAL, the effect of the clearing cannot be seen on an individual animal to animal basis, and the modest effect emerges only after group analysis. In contrast, we demonstrated dramatic improvement that was systematic across animals. We also established that our technique is compatible with calcium imaging and does not result in gross behavioral impairment.

Developments in imaging methods will also contribute to our ability to image deeper. Increasing achievable imaging depth continues to be an active area of research^49,50^. Because our technique is independent of the particular imaging hardware, we believe our clearing approach can act in a complementary manner with these deep imaging techniques, and thus have a diverse effect on the field and further extend our collective understanding of biological processes.

## Methods

### Development of clearing agent

A suitable compound for a clearing agent for living tissue should be non-toxic and capable of raising the RI substantially at a low concentration in order to minimally alter the composition of ECF. For wavelengths of visible light, RI of a non-magnetic substance can be approximated as the square root of relative permittivity^51^. The Clausius-Mossotti equation relates permittivity to molecular polarizability, with elements that are more polarizable having higher refractive index values Generally, larger atoms have greater polarizability since their electrons are more numerous and further from the nucleus. However, many heavy elements are known to be neurotoxic and are unsuitable for our purposes.

Iodine-based compounds are commonly used as intravenous radiocontrast agents since the absorption cross section increases as a function of the third power of the atomic number. Due to its utility in medical x-ray imaging, there has been focus on creating minimally-toxic iodine-based compounds. Specifically, since side effects arise from injecting hyperosmotic solutions into patients, research has gone into generating compounds with multiple iodine atoms per molecule so that fewer molecules have to be introduced for the same degree of contrast^52^. Similarly, this molecule will increase refractive index with a smaller effect on osmolarity.

Iodixanol (trade name: Visipaque 320, GE Heathcare) is a hydrophilic, non-ionic, iodinated dimer with six iodine atoms per molecule that is used clinically as a radiocontrast agent. The main advantage of Visipaque is that with an osmolality of 290 mOsm, it is isotonic to plasma, thereby reducing toxicity^53^. Since it is non-ionic, the iodine does not dissociate into ions but instead remains covalently bound to its molecular cage. Iodixanol has shown to be very stable^54,55^. The I-ACSF recipe was designed as a mixture of iodixanol and standard components of ACSF to increase the refractive index to 1.385 (184 mM iodixanol). To maintain the same osmolality as standard ACSF, the sodium chloride and glucose concentrations were reduced (see below). The refractive index values of the prepared media were measured by an Abbe refractometer. The osmolarity of the media was measured by a vapor pressure osmometer calibrated with a 290 mOsm standard (Wescor).

The cost of iodixanol can be a factor for experiments which require continuous superfusion during imaging. In addition to its clinical formulation, iodixanol can be purchased in the form of Opti-Prep (Sigma Aldrich), used as a density gradient media. An estimate of the iodixanol cost is $48 per 100 ml of I-ACSF with a RI of 1.385.

### Transmission imaging of tissue

C57BL/6 (Jackson Laboratory, RRID:IMSR_JAX:000664) and GAD65-GFP adult mice were used. All experimental procedures involving animals were approved by the Washington University Animal Studies Committee.

The following solutions were prepared: ACSF, containing NaCl 125 mM, KCl 2.5 mM, CaCl_2_ 2 mM, MgCl_2_ 10 mM, NaHCO_3_ 25 mM, NaH_2_PO_4_ 1.25 mM and glucose 25 mM; I-ACSF, containing the same concentrations of all compounds except NaCl 45 mM, 0.184 mM iodixanol and glucose 10 mM. Both media were equilibrated by bubbling with 95% O_2_/5% CO_2_.

To collect brains for slicing, the mice were euthanized by CO_2_. After decapitation, the brain was rapidly removed from the skull and sliced with a vibrating microtome (Leica VT1200S) while submerged in ice cold ACSF. The slices were incubated in ACSF at room temperature before imaging.

The slices were transferred into a chamber and placed on a resolution target (1951 USAF, Edmund Optics). The slices were superfused with either ACSF or I-ACSF while held in place with a harp. The slices were imaged with brightfield transmitted illumination using a Leica MZ75 using a 1x air objective (Plan NA 0.082, Leica) or an Olympus BX61 using a 10x water immersion objective (UMPLFL NA 0.3, Olympus). Light transmission through tissue overlaid on an opaque edge on the resolution target was imaged. The mean intensity in the direction orthogonal to the transparent-opaque boundary was calculated to estimate an edge spread function. The data was low pass filtered by applying a 61-point, cubic Savitzky-Golay filter along the axis perpendicular to the resolution target edge and differentiated numerically to obtain the approximation of the line spread function. The one-dimensional fast Fourier transform of the line spread function was plotted against spatial frequency to obtain the MTF curve.

### Stereotactic surgery and infusion

C57BL/6 adult male and female mice (Jackson Laboratory), 9-12 weeks old, were used for behavioral studies. Acetazolamide (Sigma Aldrich) was added to Visipaque (320 mg I/ml) or ACSF to form a 0.1 mM AZ solution. An osmotic pump (model 2001D, 8 µl/h, Alzet Corporation) was filled and attached via vinyl catheter tubing to a guide cannula (Plastics One). The pump-cannula setup was primed in phosphate buffered saline for 3 hours at 37°C before surgery.

Mice were anesthetized with an intraperitoneal injection of a ketamine xylazine cocktail (100 mg/kg ketamine, 10 mg/kg xylazine) and given carprofen (5 mg/kg, sc) and dexamethasone (2 mg/kg, ip). The mouse was mounted onto a stereotactic stage (Kopf Instruments) and given isoflurane (1-2%), and a midline incision was made to expose the skull. The head was leveled and the cannula was placed in the right lateral ventricle (−0.4 mm posterior to bregma, +1.0 mm lateral to the midline, −2.5 mm ventral to the skull surface) and secured with dental cement. The pump, connected to the cannula with catheter tubing, was implanted subcutaneously in the back of the mouse. For infusions longer than one day, the pumps were replaced every 24 hours.

### CT imaging

Static CT imaging was performed by the Washington University Small Animal Imaging Facility. Visipaque and ACSF infused mice were imaged in a small animal Inveon PET/CT scanner (Siemens Preclinical Solutions), placed nose to nose along the bore of the scanner under 2% isoflurane anesthesia. Cannulas made of PEEK tubing were used for all mice undergoing CT imaging to minimize beam hardening artifacts. Images were reconstructed at 50×50×90 µm voxel size. Calibration standards (0, 4.2, 21, 42, 63, 105 µM iodixanol) were imaged using the same imaging protocol. To determine tissue iodine concentration, we registered CT image stacks from each animal to a template stack then quantified the radiodensity of brain regions. We excluded the ventricles in this measurement to limit the quantification to iodixanol content in parenchyma. Data analysis was performed with Matlab (Mathworks, RRID:SCR_001622) and ImageJ (NIH, RRID:SCR_003070).

### Behavioral testing

Directly following CT imaging, and at least 30 minutes after the end of isoflurane exposure, mice underwent behavioral testing by the Washington University Animal Behavior Core. This sequence of events allowed us to limit the tested mice to those that were confirmed as being infused with iodixanol. Three groups of mice (n_non-infused_ = 12, n_ACSF-infused_ = 10, n_I-ACSF-infused_ = 10) were tested for any gross differences in behavior by a blinded observer. Non-infused B6 mice were tested to observe any change in behavior related to the surgery, the implantation of the cannula and the pump, or the infusion. The behavior tests included in this battery were performed as described previously^56,57^.

### Multielectrode recordings of the retina

Postnatal day 5 to 7 (P5-7) mice were dark-adapted (>1 hr), anesthetized with CO_2_, decapitated, and their retinas isolated under infrared illumination. We recorded large ensembles of retinal ganglion cells on planar arrays of 252 electrodes arranged in a 16 x 16 grid with the corner positions empty (electrode size: 30 μm, electrode spacing: 100 μm, Multi Channel Systems). Spontaneous activity was recorded in darkness from retinas perfused with warm (30-33° C) ACSF, I-ACSF, or ACSF with 2% isoflurane at 5-7 mL min^-1^. Signals of each electrode were filtered (300-3,000 Hz) and digitized at 10 kHz. Signal cut-outs from 1 ms before to 2 ms after crossings of negative thresholds were recorded to hard disk together with the time of threshold crossing (i.e., the spike time). We sorted spikes into trains representing the activity of individual neurons by principal component analysis of spike waveforms (Offline Sorter, Plexon, RRID:SCR_000012). When the activity of a single neuron was recorded on more than one electrode (identified by cross-correlation), we retained only the train with the most spikes in our subsequent analysis.

Spontaneous bursts of activity were identified for single units as events of >4 consecutive spikes separated by interspike intervals of <0.4 s if the Poisson probability of the respective spike train was <10^-4^. Single neuron bursts were identified as parts of retinal waves if the Poisson probability of spike trains across all other units given the number of units and their average firing rates was <10^-4^. Spike time tiling coefficients (STTCs) were computed as described by Cutts and Eglen (2014).

### Confocal imaging of live tissue

Confocal imaging was performed using an Olympus FV500 with a 20x multi-immersion objective (HC PL APO IMM CORR CS NA 0.7, Leica) and an argon-ion 488 nm laser for excitation. 200 µm coronal slices from adult mouse brains with a subset of neurons labelled with GFP (Gad65-GFP) were prepared in a manner similar as to what was described for transmission imaging. The slices were placed in an imaging chamber consisting of an o-ring attached to a resolution target. For a comparison of clearing, we imaged the same region of the same tissue in ACSF and I-ACSF by gradually replacing the media while the tissue remained under the objective. To maintain tissue health, ACSF and I-ACSF were continuously bubbled with humidified 95% O_2_/5% CO_2_, in-line filtered and constantly recirculated during imaging. The focal plane was corrected after the objective correction collar was adjusted as appropriate for the media. Stacks were acquired after waiting 30 minutes after changing between media. Stacks were imaged with 1.5 µm axial step size. Due to concerns about evaporation increasing osmolarity during imaging, samples of solution from the imaging chamber were collected throughout the session to monitor osmolarity. Osmolarity of the media was stable and remained under 320 mOsm throughout imaging. Transmission images using the illumination wavelength as used for fluorophore excitation were also obtained after fluorescence imaging in ACSF and I-ACSF, respectively, for direct comparison of clearing as measured by the two light microscopy techniques. If noted, maximum intensity projections are shown to minimize the effect of focal plane shifts due to changing the immersion media refractive index and adjusting the objective for spherical aberration. Images are displayed after a square root transform to compress the dynamic range of intensities.

### Two photon imaging

Imaging was performed with a Bruker multiphoton microscope equipped with a 10x multi-immersion objective (XLPLN10XSVMP, 0.7 NA, Olympus). Excitation was provided by a Ti:Sapphire laser (MaiTai and MaiTai DeepSee, Spectra Physics) tuned to 920 nm for GFP and GCaMP and 1040 nm for tdTomato. Live slice two photon imaging was performed as described above for confocal imaging.

### In vivo imaging

Gad65-GFP mice were used for anatomical imaging. Under isoflurane anesthesia, a cranial window (3 mm) was made in the skull, taking care to keep the dura intact. Dexamethasone (sc, 2 mg/kg) was administered to reduce inflammation, and carprofen (sc, 5 mg/kg) or buprenorphine SR (sc, 1 mg/kg) was administered for analgesia. Titanium head bars were attached to the skull with dental cement (C&B Metabond). Motion dampening was provided by a titanium mesh placed directly on top of the brain surface. The mesh provided light pressure to minimize axial motion while the pores in the mesh still allowed superfusion access to the brain. The brain surface was kept moist by a saline soaked Gelfoam sponge. Mice were injected with acetazolamide (ip, 50 mg/kg) to reduce cerebral spinal fluid production. For some imaging sessions, the mouse was anesthetized with a ketamine/xylazine cocktail. Mice were head-fixed under the microscope, and the brain exposed to superfusion media. Imaging was performed during superfusion with ACSF or I-ACSF.

Gad2-cre or Emx1-cre mice crossed to Ai95D mice (Jackson Laboratories) were used to perform calcium imaging. Mice were injected with AAV1.CAG.tdTomato.WPRE.SV40 (Penn Vector Core) at least three weeks prior to imaging. For functional imaging, a circular cranial window with a diameter of 5 mm was made over the cortex of mice with GCaMP6f labelled neurons driven by Emx1-cre. The window was made over the right somatosensory or visual cortex. Media delivery occurred using a peristaltic pump through inlet and outlet ports during imaging at a flow rate of 1 ml/min. Mice were head-fixed on a treadmill during imaging.

### Image analysis

Image analysis was done using Matlab (Mathworks) and ImageJ (NIH). For the calcium imaging datasets, a CNMF package was used to segment neurons and identify matching neurons across conditions^58^.

## Acknowledgements

This work was supported by NIH R01 DC010381 and R01 NS068409 (TEH) and R01 EY026978, R01 EY034001, and R01 EY027411 (DK).

## Author contributions

TEH conceived and supervised project. NK and MK developed clearing protocol. MK carried out imaging experiments and analysis. AA and DK performed retinal recordings and analysis of MEA data. DW and JD conducted behavioral testing and analysis; MK also did analysis of behavioral data. DWK analyzed calcium imaging data. TEH and MK wrote the manuscript.

## References

1 Tuchin, V. V. et al. Light propagation in tissues with controlled optical properties. J Biomed Opt 2, 401–417 (1997).

2 Wang, C. et al. Multiplexed aberration measurement for deep tissue imaging in vivo. Nat Methods 11, 1037–1040 (2014).

3 Wang, K. et al. Rapid adaptive optical recovery of optimal resolution over large volumes. Nat Methods 11, 625–628 (2014).

4 Ji, N. The practical and fundamental limits of optical imaging in mammalian brains. Neuron 83, 1242–1245 (2014).

5 Tuchin, V. V. Optical immersion as a new tool for controlling the optical properties of tissues and blood. Laser Phys 15, 1109–1136 (2005).

6 Bereiterhahn, J., Fox, C. H. & Thorell, B. Quantitative Reflection Contrast Microscopy of Living Cells. J Cell Biol 82, 767–779 (1979).

7 Wang, R. K. & Tuchin, V. V. Optical coherence tomography: light scattering and imaging enhancement. Handbook of coherent-domain optical methods, 665 (2013).

8 Tuchin, V. V. Tissue optics and photonics: light-tissue interaction. Journal of Biomedical Photonics & Engineering 1, 98–134 (2015).

9 Zhu, D., Larin, K. V., Luo, Q. & Tuchin, V. V. Recent progress in tissue optical clearing. Laser Photon Rev 7, 732–757 (2013).

10 Richardson, D. S. & Lichtman, J. W. Clarifying Tissue Clearing. Cell 162, 246–257 (2015).

11 Becker, K., Jahrling, N., Saghafi, S., Weiler, R. & Dodt, H. U. Chemical clearing and dehydration of GFP expressing mouse brains. PLoS One 7, e33916 (2012).

12 Chung, K. et al. Structural and molecular interrogation of intact biological systems. Nature 497, 332–337 (2013).

13 Dodt, H. U. et al. Ultramicroscopy: three-dimensional visualization of neuronal networks in the whole mouse brain. Nat Methods 4, 331–336 (2007).

14 Erturk, A. et al. Three-dimensional imaging of the unsectioned adult spinal cord to assess axon regeneration and glial responses after injury. Nat Med 18, 166–171 (2011).

15 Hama, H. et al. Scale: a chemical approach for fluorescence imaging and reconstruction of transparent mouse brain. Nat Neurosci 14, 1481–1488 (2011).

16 Ke, M. T., Fujimoto, S. & Imai, T. SeeDB: a simple and morphology-preserving optical clearing agent for neuronal circuit reconstruction. Nat Neurosci 16, 1154–1161 (2013).

17 Kuwajima, T. et al. ClearT: a detergent- and solvent-free clearing method for neuronal and non-neuronal tissue. Development 140, 1364–1368 (2013).

18 Tainaka, K. et al. Whole-body imaging with single-cell resolution by tissue decolorization. Cell 159, 911–924 (2014).

19 Yang, B. et al. Single-cell phenotyping within transparent intact tissue through whole-body clearing. Cell 158, 945–958 (2014).

20 Aoyagi, Y., Kawakami, R., Osanai, H., Hibi, T. & Nemoto, T. A rapid optical clearing protocol using 2,2’-thiodiethanol for microscopic observation of fixed mouse brain. PLoS One 10, e0116280 (2015).

21 Hama, H. et al. ScaleS: an optical clearing palette for biological imaging. Nat Neurosci 18, 1518–1529 (2015).

22 Costantini, I. et al. A versatile clearing agent for multi-modal brain imaging. Sci Rep 5, 9808 (2015).

23 Lee, E. et al. ACT-PRESTO: Rapid and consistent tissue clearing and labeling method for 3-dimensional (3D) imaging. Sci Rep 6, 18631 (2016).

24 Susaki, E. A. & Ueda, H. R. Whole-body and Whole-Organ Clearing and Imaging Techniques with Single-Cell Resolution: Toward Organism-Level Systems Biology in Mammals. Cell Chem Biol 23, 137–157 (2016).

25 Murray, E. et al. Simple, Scalable Proteomic Imaging for High-Dimensional Profiling of Intact Systems. Cell 163, 1500–1514 (2015).

26 Iijima, K., Oshima, T., Kawakami, R. & Nemoto, T. Optical clearing of living brains with MAGICAL to extend i n vivo imaging. iScience 24, 101888 (2021).

27 Ou, Z. et al. Achieving optical transparency in live animals with absorbing molecules. Science 385, eadm6869 (2024).

28 Bykov, A. et al. Imaging of subchondral bone by optical coherence tomography upon optical clearing of articular cartilage. J Biophotonics 9, 270–275 (2016).

29 Ke, M. T. et al. Super-Resolution Mapping of Neuronal Circuitry With an Index-Optimized Clearing Agent. Cell Rep 14, 2718–2732 (2016).

30 Eivindvik, K. & Sjogren, C. E. Physicochemical properties of iodixanol. Acta Radiol Suppl 399, 32–38 (1995).

31 Boothe, T. et al. A tunable refractive index matching medium for live imaging cells, tissues and model organisms. Elife 6 (2017).

32 Jacques, S. L. Optical properties of biological tissues: a review. Phys Med Biol 58, R37–61 (2013).

33 Helmchen, F. & Denk, W. Deep tissue two-photon microscopy. Nat Methods 2, 932–940 (2005).

34 Oheim, M., Beaurepaire, E., Chaigneau, E., Mertz, J. & Charpak, S. Two-photon microscopy in brain tissue: parameters influencing the imaging depth. J Neurosci Methods 111, 29–37 (2001).

35 Rudick, R. A., Zirretta, D. K. & Herndon, R. M. Clearance of albumin from mouse subarachnoid space: a measure of CSF bulk flow. J Neurosci Methods 6, 253–259 (1982).

36 Oshio, K., Watanabe, H., Song, Y., Verkman, A. S. & Manley, G. T. Reduced cerebrospinal fluid production and intracranial pressure in mice lacking choroid plexus water channel Aquaporin-1. FASEB J 19, 76–78 (2005).

37 Sykova, E. & Nicholson, C. Diffusion in brain extracellular space. Physiol Rev 88, 1277–1340 (2008).

38 Hrabetova, S. & Nicholson, C. Contribution of dead-space microdomains to tortuosity of brain extracellular space. Neurochem Int 45, 467–477 (2004).

39 Wong, R. O. Retinal waves and visual system development. Annu Rev Neurosci 22, 29–47 (1999).

40 Lopez-Bendito, G. et al. Preferential origin and layer destination of GAD65-GFP cortical interneurons. Cereb Cortex 14, 1122–1133 (2004).

41 Theer, P., Hasan, M. T. & Denk, W. Two-photon imaging to a depth of 1000 microm in living brains by use of a Ti:Al2O3 regenerative amplifier. Opt Lett 28, 1022–1024 (2003).

42 Bashkatov, A. N. et al. in *Saratov Fall Meeting 2005: Optical Technologies in Biophysics and Medicine VII*. 338–349 (SPIE).

43 Zhu, D., Wang, J., Zhi, Z., Wen, X. & Luo, Q. Imaging dermal blood flow through the intact rat skin with an optical clearing method. J Biomed Opt 15, 026008 (2010).

44 Wen, X., Mao, Z., Han, Z., Tuchin, V. V. & Zhu, D. In vivo skin optical clearing by glycerol solutions: mechanism. J Biophotonics 3, 44–52 (2010).

45 Wang, J., Shi, R. & Zhu, D. Switchable skin window induced by optical clearing method for dermal blood flow imaging. J Biomed Opt 18, 061209 (2013).

46 Wang, J., Zhang, Y., Xu, T., Luo, Q. & Zhu, D. An innovative transparent cranial window based on skull optical clearing. Laser Physics Letters 9, 469–473 (2012).

47 Zhao, Y.-J. et al. Skull optical clearing window for in vivo imaging of the mouse cortex at synaptic resolution. Light: Science & Applications 7, 17153–17153 (2018).

48 Magidson, V. & Khodjakov, A. Circumventing photodamage in live-cell microscopy. Methods in cell biology 114, 545–560 (2013).

49 Kondo, M., Kobayashi, K., Ohkura, M., Nakai, J. & Matsuzaki, M. Two-photon calcium imaging of the medial prefrontal cortex and hippocampus without cortical invasion. Elife 6 (2017).

50 Ouzounov, D. G. et al. In vivo three-photon imaging of activity of GCaMP6-labeled neurons deep in intact mouse brain. Nat Methods 14, 388–390 (2017).

51 Dressel, M. & Grüner, G. Electrodynamics of solids: optical properties of electrons in matter. (Cambridge university press, 2002).

52 Hogstrom, B. & Ikei, N. Physicochemical properties of radiographic contrast media, potential nephrotoxicity and prophylaxis. Clinical and Experimental Pharmacology and Physiology 42, 1251–1257 (2015).

53 Karlsson, J., Gregersen, M. & Refsum, H. Visipaque is isotonic to human and rat blood plasma. Acta Radiologica 36, 39–42 (1995).

54 Krause, W. & Schneider, P. W. in *Contrast agents II: Optical, ultrasound*, X-ray and radiopharmaceutical imaging 107–150 (Springer, 2002).

55 Priebe, H., Aukrust, A., Bjørsvik, H. R., Tønseth, C. & Wiggen, U. Stability of the X-ray contrast agent iodixanol= 3, 3′, 5, 5′-tetrakis (2, 3-dihydroxypropylcarbamoyl)-2, 2′, 4, 4′, 6, 6′-hexaiodo-N, N′-(2-hydroxypropane-1, 3-diyl)-diacetanilide towards acid, base, oxygen, heat and light. Journal of clinical pharmacy and therapeutics 24, 227–235 (1999).

56 Wozniak, D. F. et al. Motivational disturbances and effects of L-dopa administration in neurofibromatosis-1 model mice. PLoS One 8, e66024 (2013).

57 Barnes, T. D. et al. A Mutation Associated with Stuttering Alters Mouse Pup Ultrasonic Vocalizations. Curr Biol (2016).

58 Giovannucci, A. et al. CaImAn an open source tool for scalable calcium imaging data analysis. Elife 8 (2019).

